# Mirror-image inversion in commonly used compound microscopes

**DOI:** 10.1101/2025.09.04.674179

**Authors:** François Lapraz, Céline Boutres, Baptiste Monterroso, Stéphane Noselli

## Abstract

Optical microscopes are essential tools for magnifying and analyzing a wide range of samples and are expected to faithfully preserve the original object in all its dimensions. However, while examining cerebral structures with left–right asymmetry, we discovered that the image of the object can be unintentionally mirror-inverted, depending on the microscope setup. This inversion mostly goes unnoticed, leading to incorrectly oriented images, loss of image integrity, and potential misinterpretation. We detail the optical origins of this issue and offer a practical guide to support accurate sample imaging, while urging manufacturers to explicitly communicate how their systems affect image orientation.

## MAIN TEXT

Compound microscopes magnify samples using a series of optical elements, each impacting light quality in specific ways. Key components include the condenser, objective, dichroic filter, prism, beamsplitter, mirror, eyepiece, and detector. These elements magnify (lens, eyepiece), correct aberrations (lens), split (dichroic, beamsplitter), reflect (mirror), or collect light (detector, eyepiece). Figure 1a shows each element’s effect on the image. Some elements have a neutral while others can have a more transformative effect. For instance, lenses flip the image both vertically and horizontally, but a simple 180° rotation in the lens plane restores the original orientation without altering its three-dimensional structure (Figure 1b). However, reflective elements like mirrors, dichroics, and beamsplitters are more concerning as they create mirror images, changing the object’s 3D-structure (Figure 1b). To illustrate the significance of this transformation, consider our hands: their 3D-structures are mirror images and cannot be superimposed, highlighting the concept of chirality. This property is prevalent in nature, influencing biological molecules (L- and D-amino acids, L- and D-sugars, F-actin, microtubules), cells (e.g., polarized and asymmetrically dividing cells), tissues/organ shape (e.g., plant twining) and position (e.g., heart, gut), and the entire organism (e.g., spiraling snail shell). Any chiral object (e.g., a right hand) will retain its 3D structure when viewed through a lens; however, upon reflection in a mirror, it will appear in its opposite chiral form (e.g., as a left hand) (Figure 2b).

**Figure 1:**
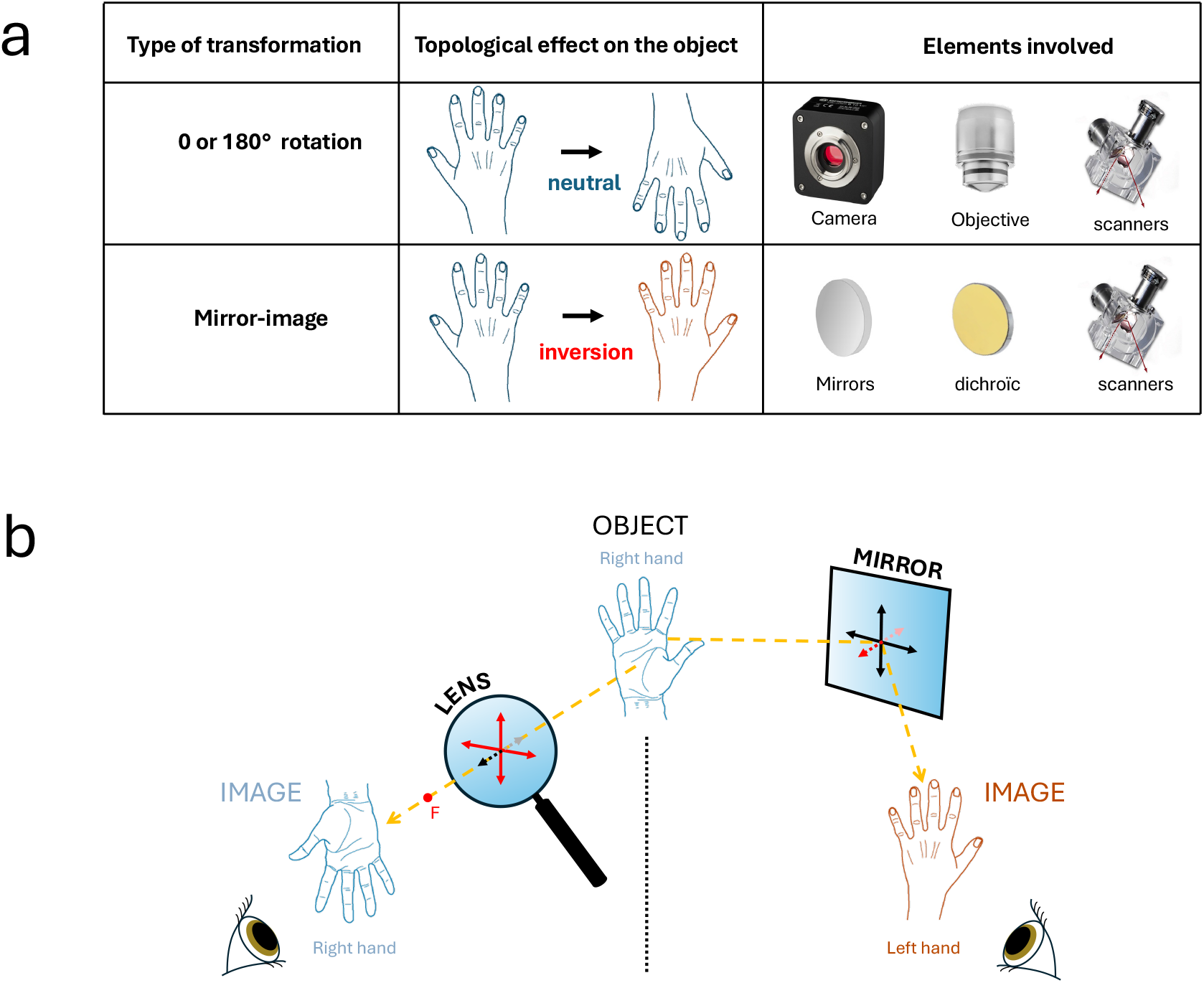
Image transformations caused by standard microscope elements. **a)** 2-Dimensionnal geometric transformations of an image caused by common optical and mechanical elements. Each element in the system - transmission, detection, and sopware - combine to produce the final image. The lep and right hands are used as chiral objects to best illustrate the transformations. **b)** Images observed through a convergent lens aper its focal point (F) are inverted along the axes parallel to the plane of the lens (Red arrows). Right hand observed under these conditions produces an image of a right hand with a 180° rotation in the lens plane. Images observed aper reflection in a plane mirror are inverted along the axis perpendicular to the plane of the mirror (Red arrows). A right hand observed under these conditions produces an image of a lep hand.

**Figure 2:**
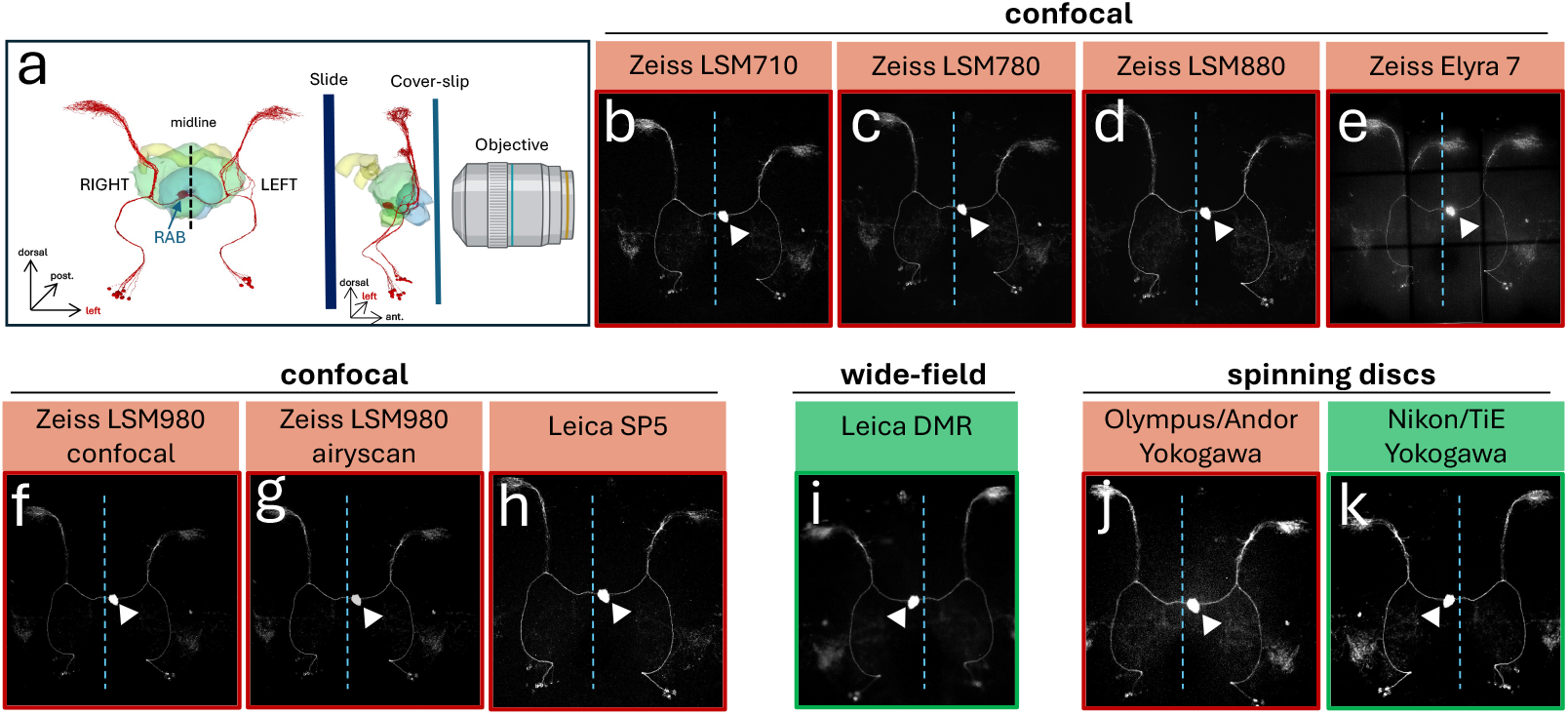
The visualization of asymmetric H-neurons under various microscope setups. **a)** Schematic representation of H-neurons (red) and the central complex of the adult brain (yellow, green, blue). The H-neurons project their axons into the right neuropile (right asymmetric body, RAB; arrow). In a frontal view, the RAB appears on the lep side. The midline is indicated by a dashed line. Single brain preparations were imaged with the objective facing the anterior side (frontal view-right panel). In this setup, the resulting image should display the RAB on the lep side, as shown in the lep panel. **b-k)** Images of the same preparation (*72A10-LexA, LexAop-CD4-tdTomato* fly brain; mounted as in **a)**, were captured using various microscope setups (confocal, spinning discs, wide field) from different manufacturers. Depending on the setup, images are either inverted (b-h, j) (red outline) or in the correct orientation **(i, k)** (green outline). The midline is indicated by a blue dashed line. RAB is indicated by an arrowhead.

While imaging a left-right (LR) asymmetric neuronal circuit with different microscope setups, we discovered that, depending on the setup used, the output image could be either in its original or mirror-imaged conformation. In adult Drosophila, H-neurons form a small asymmetric circuit with axons projecting specifically to the right asymmetrical body (RAB) in the brain [1,2]. Imaging these neurons on various systems, we found that some produced accurate images (neurons projecting to the RAB), while others displayed a mirror-image object, showing projections to the left AB (Figure 2). Without clear LR markers like H-neurons, this inversion would have gone unnoticed. If these cells bore text on their surfaces, it would appear inverted, like a reflection in a mirror.

Discussions with colleagues and imaging platform heads revealed that most users are unaware of this image transformation issue. Our sampling included seven inverted microscope systems from three different brands and two upright systems from a distinct fourth brand (Figure 2; Table 1), indicating that the inversion problem is not confined to a single manufacturer or product line. Among inverted systems, six showed mirror-image inversion at both eyepiece and detector, while one displayed discordance between the two. The two upright systems showed correct orientation at the eyepiece, but one of them was discordant at the detector. Thus, while all inverted microscopes tested produced inverted images at the eyepiece and all tested upright systems preserved orientation at the eyepiece, the presence in our panel of two systems with discordant eyepiece/detector orientations prevents drawing a strict divide between the two configurations, adding to the confusion (Table 1). Expanding the range of instruments tested will help confirm these trends.

**Table 1:** Summary of image transformations observed under various microscope setups at the detector and eyepiece.

|  | Brand | Model | eyepiece | detector |
| --- | --- | --- | --- | --- |
| INVERTED | Zeiss | LSM710 | inverted | inverted |
|  |  | LSM780 | inverted | inverted |
|  |  | LSM880 | inverted | inverted |
|  |  | LSM980 | inverted | inverted |
|  |  | Elyra 7 | inverted | inverted |
|  | Olympus | Spinning discs | inverted | inverted |
|  | Nikon | Spinning discs | inverted | correct |
| UP-RIGHT | Leica | SP5 | correct | inverted |
|  |  | DMR | correct | correct |

What causes image inversion, and why do some microscopes invert images while others don’t? Optical elements that simply transmit light (lens, eyepiece) do not alter the image. In contrast, reflective elements (dichroic, mirror) produce a mirror image, leading to inversion (see Figure 1b). Notably, inversion induced by reflection can be reversed by a second reflection. Therefore, the number of reflectors in the emission light path determines the final image’s orientation: if the number of reflective elements is even, the image remains true to the original; if odd, the image is inverted (Figure 3).

**Figure 3:**
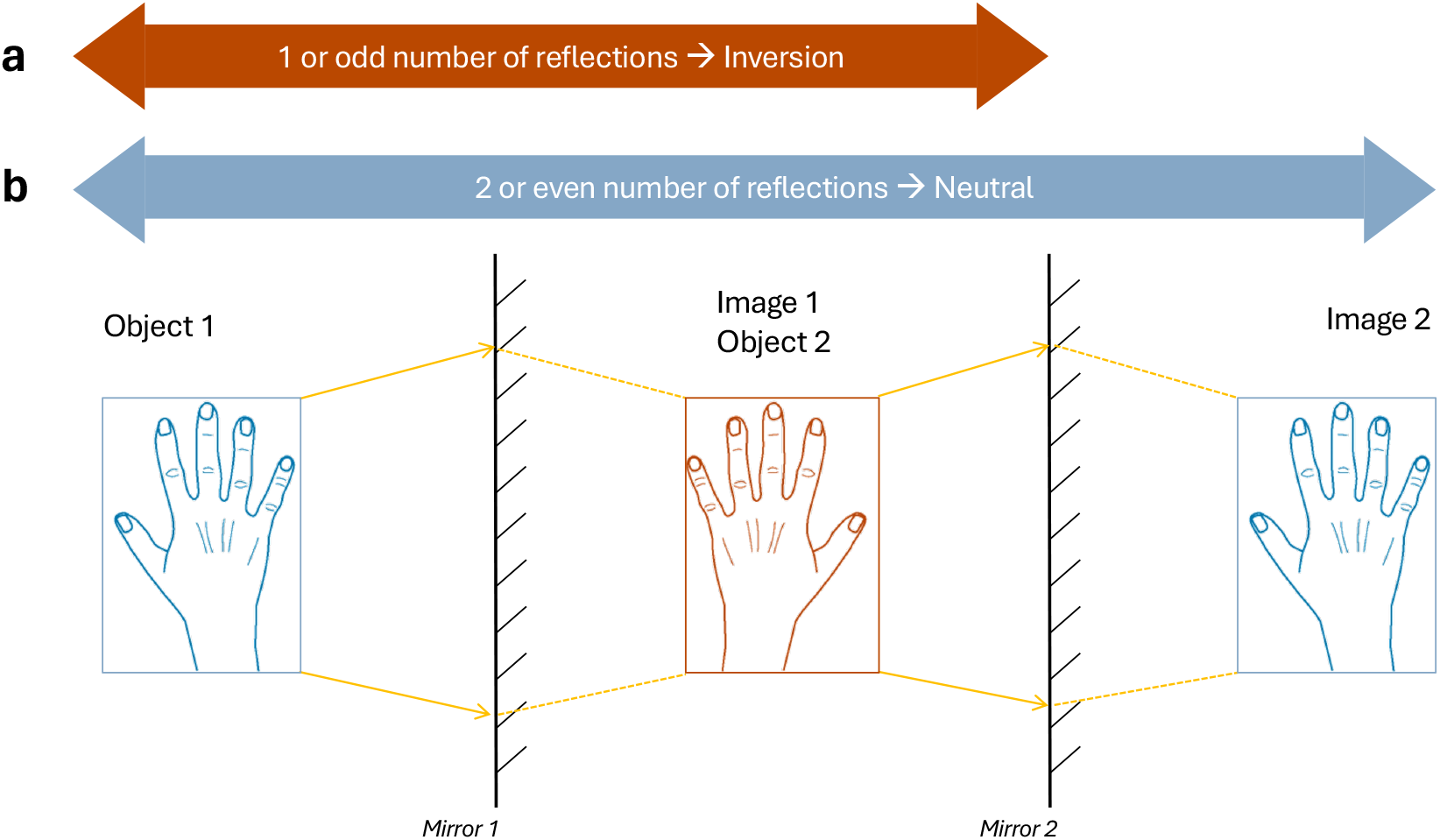
Impact of an odd or even number of reflective elements on an image. **a)** One or an odd number of mirrors: an object’s image undergoes lep/right inversion, a characteristic known as reversibility in a plane mirror. **b)** Double or an even number of mirrors: the object’s image undergoes lep/right inversion in the first mirror and is then inverted again by the second mirror. A system with mirror parity retains the primary object’s3-D structure in its final image.

Due to the mirror-image inversion generated by some equipment, many published images may actually be mirror images of the objects they describe. Without being able to trace the original setup, these images become unsuitable for future use. This issue is critical for samples with chirality, potentially altering conclusions.

We want to highlight the broader issue of mirror image transformation. Image inversion can also easily happen during post-imaging manipulation. Common software tools, like Fiji [3], have functions such as ‘Flip’ (horizontal, vertical, Z-axis), which create a mirror image when applied to an image. Consequently, any ‘Flip’ inversion along one (or an odd number) of the three spatial axes can inadvertently produce a mirror image, compromising data accuracy. Finally, image inversion can also occur during acquisition [4,5], for example, if the sensor acquires pixel columns from right to left but they are displayed from left to right in the acquisition software.

To tackle this critical yet overlooked issue, we strongly urge manufacturers to update their information sheets and inform users about the impact of their equipment on image orientation at both ocular and detector sites. Ideally, sopware should automatically correct orientation at the detector site, eliminating the need for user intervention. Furthermore, published images should be clearly labelled with their 3D orientation, including the LR axis, to ensure accurate interpretation and reproducibility. Implementing dedicated, asymmetric control slides, e.g. stage micrometers with characters on them (text and/or number), in imaging facilities is essential to resolve this issue and prevent future errors.

## METHODS

### Drosophila strains and imaging

H-neurons were visualized in flies expressing tdTomato under the control of the H-neuron-specific 72A10-LexA line (*72A10-LexA; 13xLexAop2-CD4-tdTomato*), as described in [2]. LexA and LexAop lines were obtained from the Bloomington *Drosophila* Stock Center (BDSC, Bloomington, IN, USA): 72A10-LexA (#54191); 13xLexAop2-CD4-tdTomato (#77139).

Imaging was performed on the same individual brain using Leica SP5 (Leica DM6000 CS upright stand with Leica TCS SP5 II confocal scanner unit mounted on the back), Zeiss LSM710 (Zeiss Axio Observer. Z1 inverted stand with Zeiss LSM 710 confocal scanner unit mounted on the side), LSM780 (Zeiss Axio Observer. Z1 inverted stand with Zeiss LSM 780 confocal scanner unit mounted on the back), LSM880 (Zeiss Axio Observer inverted stand with Zeiss LSM 880 confocal scanner unit and Airyscan detector mounted on the side), LSM980 (Zeiss Axio Observer inverted stand with Zeiss LSM 980 confocal scanner unit and Airyscan 2 detector mounted on the back or Elyra7 (Zeiss Axio Observer inverted stand with Zeiss Elyra 7 illumination and detection module mounted on the back) confocal microscopes, Olympus/Andor Yokogawa (Olympus IX81 inverted stand Yokogawa CSU-X1 Confocal Scanner Unit mounted on the side / Andor TuCam / Andor Ixon+ 897) or Nikon/TiE Yokogawa (Nikon eclipse Ti inverted stand with Yokogawa CSU-W1 Confocal Scanner Unit mounted on the side / Andor Ixon life 888) spinning-disc microscopes, and Leica DMR (upright stand with Lumenera infinity 3 camera mounted on eyepiece) widefield microscope.

## ACKNOWLEDGEMENTS

We wish to thank Rob Arkowitz, Alexander Bershadsky, Florence Besse, Max Fürthauer, Mazeo Rauzi, Pavel Tomancak for discussions. Work in SN laboratory is supported by Agence Nationale pour la Recherche (ANR-20-CE13-0004, ANR-24-CE13-1857, ANR-25-CE13-3952), Fondation pour la Recherche Médicale (FRM; EQU201903007825), Université Côte d’Azur (UniCA), Centre National pour la Recherche Scientifique (CNRS), Institut National pour la Recherche Médicale (Inserm).

## Notes

### Competing Interest Statement

The authors have declared no competing interest.

### Summary of Updates

The article has been revised to correct the author list.

## REFERENCES

1. Wolff, T. and Rubin, G.M. (2018) Neuroarchitecture of the Drosophila central complex: A catalog of nodulus and asymmetrical body neurons and a revision of the protocerebral bridge catalog. J. Comp. Neurol. 526, 2585–2611

2. Lapraz, F. et al. (2023) Asymmetric activity of NetrinB controls laterality of the Drosophila brain. Nat. Commun. 14, 1052

3. Schindelin, J. et al. (2012) Fiji: an open-source platform for biological-image analysis. Nat. Methods 9, 676–682

4. Schlegel, P. et al. (2024) Whole-brain annotation and multi-connectome cell typing of Drosophila. Nature 634, 139–152

5. Dorkenwald, S. et al. (2024) Neuronal wiring diagram of an adult brain. Nature 634, 124–138

